# Peroxisomal ATP uptake is managed by the ABC transporters and two adenine nucleotide transporters

**DOI:** 10.1101/2021.06.14.448280

**Authors:** Carlo van Roermund, Lodewijk IJlst, Nicole Linka, Ronald J.A. Wanders, Hans R. Waterham

## Abstract

Peroxisomes are essential organelles involved in various metabolic processes, including fatty acid β-oxidation. Their metabolic functions require a controlled exchange of metabolites and co-factors, including ATP across the peroxisomal membrane. We investigated which proteins are involved in the peroxisomal uptake of ATP in the yeast *Saccharomyces cerevisiae*. Using wild-type and targeted deletion strains, we measured ATP-dependent peroxisomal octanoate β-oxidation, intra-peroxisomal ATP levels employing peroxisome-targeted ATP-sensing reporter proteins, and ATP uptake in proteoliposomes prepared from purified peroxisomes. We show that intra-peroxisomal ATP levels are maintained by different peroxisomal membrane proteins each with different modes of action: (1) the previously reported Ant1p protein, which catalyzes ATP/AMP exchange (2) the ABC transporter protein complex Pxa1p/Pxa2p, which mediates both acyl-CoA and ATP uptake; and; (3) the mitochondrial Aac2p protein, which catalyzes ATP/ADP exchange and was shown to have a dual localization in both mitochondria and peroxisomes. Our results provide compelling evidence for an ingenious complementary system for the uptake of ATP in peroxisomes.

## Introduction

Peroxisomes are single-membrane bounded organelles found in cells of all eukaryotic species. They can be involved in a large variety of metabolic pathways which may differ per species but always includes the degradation of fatty acids through β-oxidation. In mammals, including humans, peroxisomes also play an important role in ether phospholipid biosynthesis, fatty acid alpha-oxidation, bile acid synthesis, glyoxylate detoxification and H_2_O_2_ degradation^1, 2^. Genetic defects in the biogenesis and/or functioning of peroxisomes affect these metabolic pathways and usually have severe clinical consequences, as is dramatically demonstrated in the Zellweger spectrum disorders^3^ and the various single peroxisomal enzyme deficiencies^4^.

In most metabolic pathways, peroxisomes only catalyze a specific subset of enzyme reactions with other reactions catalyzed in the cytosol, mitochondria and/or endoplasmic reticulum^5^. This implies that the various metabolites involved, i.e. substrates and products, and co-factors, i.e. NAD, ATP, CoA, need to be transported across the peroxisomal membrane. In the past decades, the enzymology and biochemical functions of peroxisomes have largely been resolved. However, the mechanisms involved in peroxisomal metabolite transport have remained largely unknown. Yet, the importance of this transport is underlined by the existence of two inherited human diseases that are caused by defects in the peroxisomal half ABC transporter proteins ABCD1 and ABCD3^4^, which function in the peroxisomal import of the CoA esters of very-long-chain fatty acids and branched-chain fatty acids, respectively.

Current consensus holds that peroxisomes are equipped with two fundamentally different mechanisms for metabolite transport across their membrane, which includes (1.) diffusion of small Mw metabolites (<400Da) via channel-forming membrane proteins, and (2.) carrier-mediated transport of higher Mw metabolites, such as acyl-CoAs and ATP. Genetic complementation approaches, sequence similarity searches, and proteomic analyses of highly purified peroxisomes of mouse^6, 7^, human^8^, plants^9^, and the yeast *Saccharomyces cerevisiae*^10, 11^ have led to the identification of several integral peroxisomal membrane proteins which, based on (partial) sequence similarity shared with known transport proteins, may function as peroxisomal metabolite transport proteins.

In mammals, including humans, three half ABC transporter proteins have been identified in the peroxisomal membrane: ABCD1 (also known as adrenoleukodystrophy protein, ALDP)^12, 13^, ABCD2 (also known as adrenoleukodystrophy-related protein, ALDR)^14, 15^ and ABCD3 (also known as 70-kDa peroxisomal membrane protein, PMP70)^16^. These proteins were shown to function as homodimers and import long, very-long-chain and branched-chain acyl-CoA esters into peroxisomes^17, 18^. So far, only three additional mammalian peroxisomal membrane proteins with a presumed function in metabolite transport have been identified. The first one is SLC25A17, also known as PMP34, which, based on sequence similarity, is a member of the mitochondrial carrier family (MCF). Reconstitution experiments in proteoliposomes indicated that this protein may function as a CoA/NAD/FAD transporter^19^. The second protein is PXMP2, which was shown to have channel-forming properties^20^. The third protein is PXMP4, which shares some similarity with bacterial permeases, but has not been functionally studied^21, 22^.

Peroxisomes in *S. cerevisiae* contain two half ABC transporters, Pxa1p and Pxa2p, which are involved in the import of long-chain acyl-CoA esters (e.g. C18:1)^23, 24, 25, 26^. In contrast to their human orthologues, Pxa1p and Pxa2p were shown to function as heterodimers. Two additional peroxisomal membrane proteins with a presumed function in metabolite transport have been identified in *S. cerevisiae*. Ant1p is an MCF member with strong similarity to human PMP34 but, in contrast to PMP34, shown to catalyze the exchange of cytosolic ATP for peroxisomal AMP. This AMP is generated upon the intra-peroxisomal ATP-dependent activation of fatty acids by the acyl-CoA synthetase Faa2p^27, 28^. The second protein, Pex11p, is known to be involved in peroxisomal fission, but also was reported to have transport or channel-forming properties^29, 30^.

We use *S. cerevisiae* as model system^31^ to unravel the mechanism of metabolite transport across the peroxisomal membrane. In contrast to human cells, in which both mitochondria and peroxisomes perform β-oxidation, fatty acid degradation in yeast cells takes place exclusively in peroxisomes and thus requires the import of fatty acids, and the co-factors ATP and CoA. Medium-chain fatty acids with carbon lengths of 8-12 enter yeast peroxisomes in their free acid form and are activated into CoA esters inside peroxisomes via the peroxisomal acyl-CoA synthetase Faa2p^23, 26^. This activation is ATP- and CoA-dependent. Long-chain fatty acids, however, are first activated outside peroxisomes and then imported as acyl-CoA ester by the ABC transporter protein complex Pxa1p/Pxa2p^23,26^, followed by release of coenzyme A at the luminal side of peroxisomes and re-esterification by a peroxisomal synthetase^17, 32, 33^.

Although in yeast, intra-peroxisomal ATP is essential for the peroxisomal β-oxidation of fatty acids following their import via the free fatty-acid route as well as the ABC transporter protein-mediated pathway, relatively little is known about the peroxisomal uptake of ATP except for the above mentioned involvement of Ant1p^27, 28^. Our observation that a knock-out of Ant1p in *S. cerevisiae* does not completely abolishes peroxisomal β-oxidation, however, implied the existence of additional ways to import ATP into peroxisomes. In this study we show that the uptake of ATP into peroxisomes is indeed mediated by different peroxisomal membrane proteins. In addition to Ant1p, these include the ABC transporter protein complex Pxa1p/Pxa2p, which thus catalyses peroxisomal ATP uptake as well as acyl-CoA import, and the MCF carrier Aac2p, a predominantly mitochondrial protein, which we found partially localized to peroxisomes and which catalyzes the exchange of cytosolic ATP for peroxisomal ADP.

## Results

### Ant1p and the ABC transporter protein complex Pxa1p/Pxa2p transport ATP across the peroxisomal membrane

We previously showed that in the yeast *S. cerevisiae*, medium chain fatty acids such as octanoate (C8:0) are imported into peroxisomes as free fatty acids. To become substrate for β-oxidation they subsequently are activated into their corresponding fatty acyl-CoA ester by the intra-peroxisomal ATP-dependent acyl-CoA synthetase Faa2 (Fig. 1A)^23, 26^. In accordance with this, the β-oxidation of C8:0 in mutant cells in which the *FAA2* gene is deleted (*faa2Δ*) is fully deficient, similar as in *fox1Δ* cells in which the *FOX1* gene encoding acyl-CoA oxidase, the first enzyme of the β-oxidation pathway, is deleted (Fig. 1B). Earlier work also showed that the peroxisomal membrane protein Ant1p functions as an ATP/AMP antiporter and thus most probably is responsible for the peroxisomal uptake of ATP required for the intra-peroxisomal activation of fatty acids^27, 28^. Indeed, deletion of *ANT1* resulted in a significant decrease in the C8:0 β-oxidation activity (Fig. 1B). However, the C8:0 β-oxidation activity in the *ant1*Δ cells was still ~30% of the activity measured in wild-type cells, which implied the involvement of additional ATP uptake system(s) in the peroxisomal membrane. To identify these, we measured C8:0 β-oxidation activities in *ant1*Δ cells in which in addition genes encoding other known peroxisomal membrane proteins were deleted. Surprisingly, we observed that C8:0 β-oxidation activity was further reduced to ~10% when we also deleted both *PXA1* and *PXA2* in the *ant1*Δ cells (Fig. 1B). In *pxa1*Δ *pxa2*Δ double mutant cells, the C8:0 β-oxidation was only slightly decreased compared to wild-type cells. These findings pointed to a novel, unanticipated role for the ABC transporter protein complex Pxa1p/Pxa2p in peroxisomal ATP uptake in addition to its established role in the peroxisomal import of fatty acyl-CoAs.

**Figure 1.**
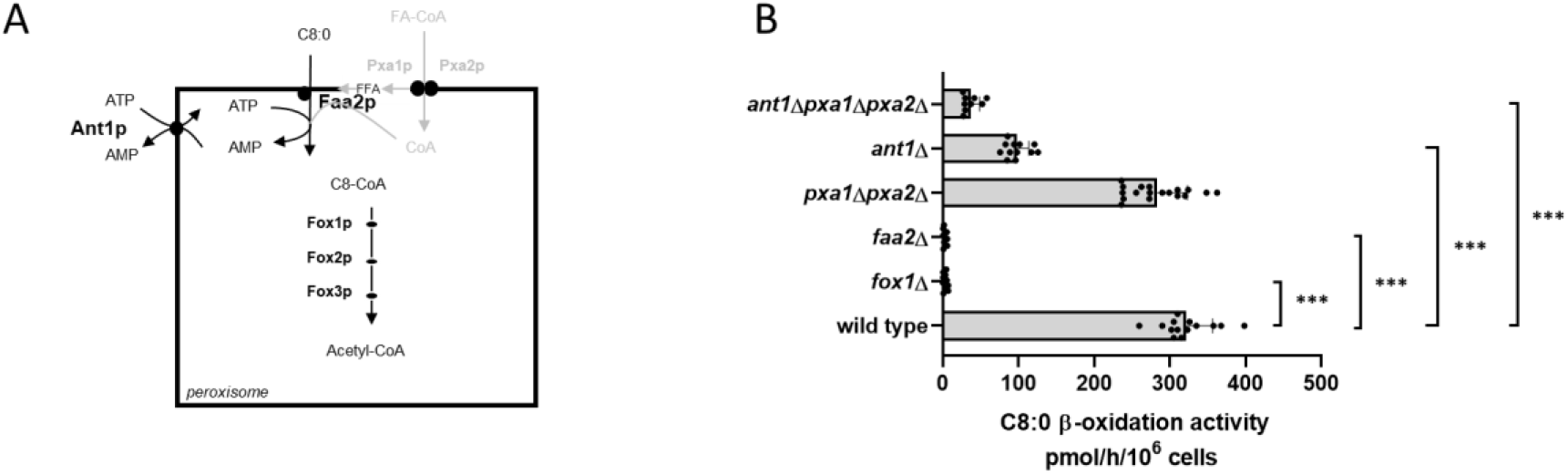
C8:0 β-oxidation activity in wild-type and mutant yeast strains. (**A**) In yeast, β-oxidation of fatty acids occurs exclusively in peroxisomes requiring the import of fatty acids, ATP and CoA. Medium-chain fatty acids (C8-C12) enter yeast peroxisomes as free fatty acids and are subsequently activated to their corresponding CoA ester by the peroxisomal enzyme acyl-CoA synthetase Faa2p. This activation step is ATP and CoA dependent. (**B**) Yeast cells were cultured overnight in oleate medium and β-oxidation rates were measured using [1-^14^C] labelled octanoate (C8:0) as substrate. Data are means ± SD of values from 5#x2013;18 independent experiments. A one-way ANOVA test with Tukey’s multiple comparisons test was performed. ***, ** and * indicate significance with a P-value of p<0.001, p<0.01 and p<0.05 respectively.

### ATP uptake in proteoliposomes prepared from peroxisomes of different mutant strains

To find additional evidence for a role of Pxa1p/Pxa2p in peroxisomal ATP import, we next studied the uptake of radio-labeled ATP in proteoliposomes prepared from peroxisomal membranes isolated from wild-type cells, *ant1*Δ, *pxa1*Δ *pxa2*Δ, and *ant1*Δ *pxa1*Δ *pxa2*Δ mutant cells.

In the absence of internal adenine nucleotides as counter-exchange substrate, we observed low and similar levels of ATP uptake in proteoliposomes prepared from *ant1Δ* and wild-type cells, while no ATP uptake was observed in proteoliposomes prepared from *pxa1*Δ *pxa2*Δ and *ant1*Δ *pxa1*Δ *pxa2*Δ mutant cells (Figure 2A).

**Figure 2.**
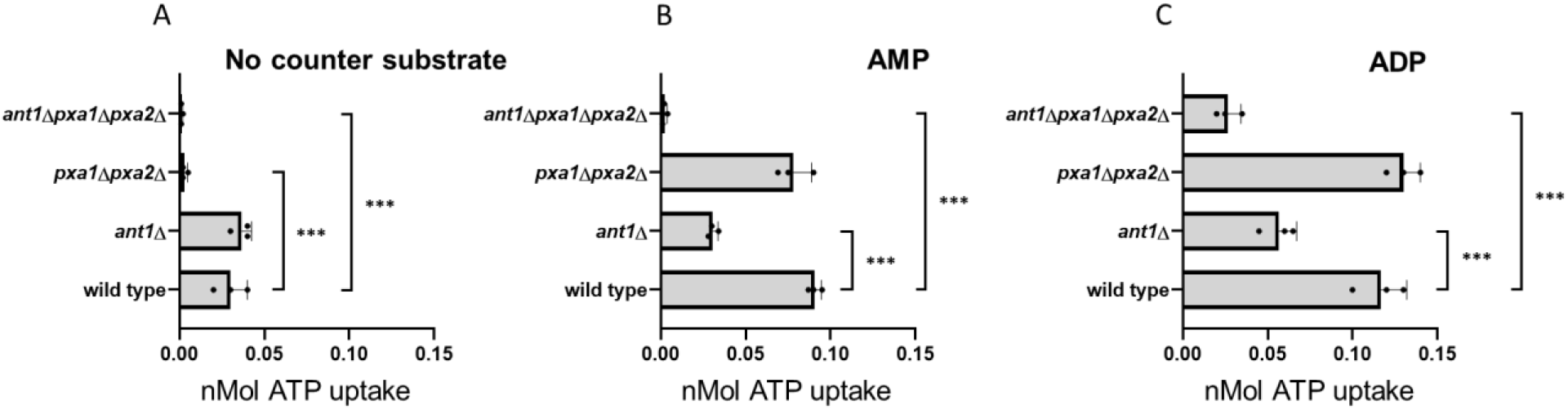
ATP uptake into liposomes reconstituted with peroxisomal membranes purified from different yeast strains. Isolated peroxisomal membranes were reconstituted in proteoliposomes with no counter substrate (**A**), or pre-loaded with AMP (**B**) or ADP (**C**). Uptake of [α-^32^P]ATP was measured as described in Methods. Data are means ± SD of values from three independent experiments. A one-way ANOVA test with Tukey’s multiple comparisons test was performed for n=3. ***, ** and * indicate significance with a P-value of p<0.001, p<0.01 and p<0.05 respectively.

When we preloaded the proteoliposomes with AMP, ATP uptake was reduced to ~30% in proteoliposomes prepared from *ant1*Δ mutant cells, slightly reduced in proteoliposomes prepared from *pxa1*Δ *pxa2*Δ mutant cells, and virtually absent in proteoliposomes prepared from *ant1*Δ *pxa1*Δ *pxa2*Δ mutant cells when compared to ATP uptake in proteoliposomes prepared from wild-type cells (Fig. 2B). These findings i) show that Ant1p is responsible for most of the peroxisomal ATP uptake, ii) confirm that Ant1p functions as peroxisomal ATP/AMP antiporter^27, 28^; iii) are in agreement with the C8:0 β-oxidation activities measured in the corresponding mutant cells; and iv) support a role for Pxa1p/Pxa2p in the peroxisomal uptake of ATP.

When we preloaded the proteoliposomes with ADP, ATP uptake was similar in proteoliposomes prepared from *pxa1*Δ *pxa2*Δ mutant cells and wild-type cells, and reduced to ~45% in proteoliposomes prepared from *ant1Δ* mutant cells. Interestingly, however, *ant1*Δ *pxa1*Δ *pxa2*Δ mutant cells still showed ~20% ATP uptake when compared to wild-type cells (Figure 2C). This confirms that Ant1p can also function as an ATP/ADP antiporter as previously shown^34^, but also indicates the involvement of at least one additional peroxisomal ATP transporter that can mediate ATP/ADP exchange.

### Direct measurement of intra-peroxisomal ATP levels

To provide *in vivo* evidence for the role of Ant1p and Pxa1/Pxa2 in peroxisomal ATP uptake, we expressed modified versions of the FRET-based ATeam reporter protein35, 36 in the different yeast strains to measure the relative ATP levels in the peroxisomes (ATeam-per, i.e. extended with a carboxy-terminal peroxisomal targeting signal) and the cytosol (ATeam-cyt). To correct for ATP-independent background fluorescence, we used mutated versions of the same reporter proteins that have no affinity for ATP (ATeam-permut and ATeam-cytmut). The relative ATP levels in cytosol and peroxisomes of *pxa1*Δ *pxa2*Δ mutant cells were similar as observed in cytosol and peroxisomes of wild-type cells (Fig. 3). However, the relative ATP levels in peroxisomes of *ant1*Δ and *ant1*Δ *pxa1*Δ *pxa2*Δ mutant cells were markedly reduced (Fig. 3A) while the relative cytosolic ATP levels remained virtually unchanged when compared to wild-type cells (Fig. 3B). The observation that the peroxisomal ATP levels in the *ant1*Δ *pxa1*Δ *pxa2*Δ mutant cells were lower than in the *ant1*Δ cells confirmed that Pxa1p/Pxa2p also mediates ATP uptake, in addition to Ant1p.

**Figure 3.**
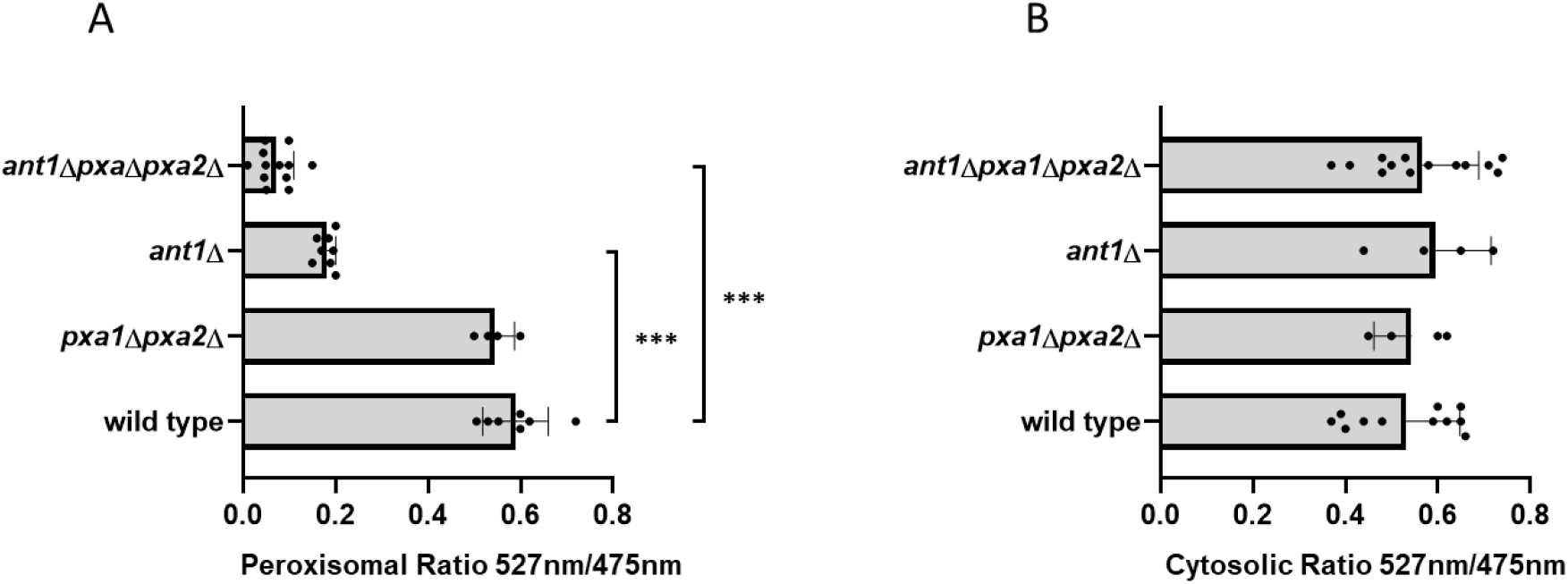
Comparison of ATP levels determined with FRET-based ATeam reporter proteins in wild-type and mutant yeast strains. Cells were transformed with the different ATeam reporter proteins, grown overnight in oleate medium and transferred to 25mM MES buffer (pH 6.0) supplemented with 20g/L glucose prior to measurements. (**A**) Peroxisomal ATP levels determined with reporter proteins ATeam-per and ATeam-per(mut) (**B**) ATP levels determined with reporter proteins ATeam-cyt and ATeam-cyt(mut). FRET was determined by measuring 527/475nm. Ratios were calculated after subtraction of the background signal from wild-type cells transformed with an empty vector. The semi-relative ATP levels were then calculated by subtracting the 572/475 ratio of the ATeam(mut) reporter from the 572/475 of ATeam reporter. Data are means ± SD of values from 4–13 independent experiments. A one-way ANOVA test with Tukey’s multiple comparisons test was performed. ***, ** and * indicate significance with a P-value of p<0.001, p<0.01 and p<0.05 respectively.

### Identification of an additional peroxisomal ATP transporter

The ~10% residual C8:0 β-oxidation activity measured in the *ant1*Δ *pxa1*Δ *pxa2*Δ mutant cells (Fig. 1B) as well as the ~20% residual *in vitro* ATP/ADP exchange observed in proteoliposomes prepared from peroxisomes of the *ant1*Δ *pxa1*Δ *pxa2*Δ mutant cells (Fig. 2C), indicated the involvement of at least one additional peroxisomal ATP transporter. We hypothesized that in addition to Ant1p, which is a member of the MCF family and exclusively localized to peroxisomes, there might be one or more other members of this carrier family that have a dual localization in both the mitochondrial and the peroxisomal membrane. In order to study this, we developed a sensitive cell-based assay employing self-assembling GFP^37^ that allows to determine a possible peroxisomal localization of MCF proteins (see Fig. S5 and S6). To this end, we tagged the C-terminus of selected MCF proteins with the GFP(11) peptide sequence, and co-expressed these with a peroxisome-targeted GFP(1-10)-PTS1 in wild-type yeast cells that also express a peroxisomal RFP-PTS1 reporter protein. We tested three mitochondrial MCF proteins that are known to function as adenine nucleotide carriers: Aac2p, Yea6p and Leu5p. As positive control we used Ant1p, and as negative control we used the mitochondrial MCF protein Sco2p, which is known to mediate copper transport to cytochrome c oxidase and thus is assumed not to have a dual localization in peroxisomes.

In contrast to GFP(11)-tagged Ant1p, we did not observe GFP fluorescence for GFP(11)-tagged Yea6p, Leu5p and Sco2p when co-expressed with peroxisome-targeted GFP(1-10)-PTS1 (Fig. 4). This implies that, like Sco2p, Yea6p and Leu5p are not co-localized to peroxisomes. GFP(11)-tagged Aac2p, however, displayed a clear punctated GFP fluorescence pattern similar as observed for GFP(11)-tagged Ant1p, which indicates a peroxisomal co-localization. This is confirmed by the co-localization of the punctated green GFP fluorescence with the peroxisomal red RFP-SKL fluorescence (Fig. 4). These findings imply that Aac2p has a dual subcellular localization in both mitochondria and peroxisomes and thus could be responsible for the residual peroxisomal ATP in peroxisomes observed in the *ant1*Δ *pxa1*Δ *pxa2*Δ mutant cells. Interestingly, Aac2p was shown previously to function as an ATP/ADP carrier ^34, 38,39, 40^, which fits very well with the ressidual *in vitro* ATP/ADP exchange we measured in proteoliposomes prepared from peroxisomes of the *ant1*Δ *pxa1*Δ *pxa2*Δ mutant cells (Fig. 2C).

**Figure 4.**
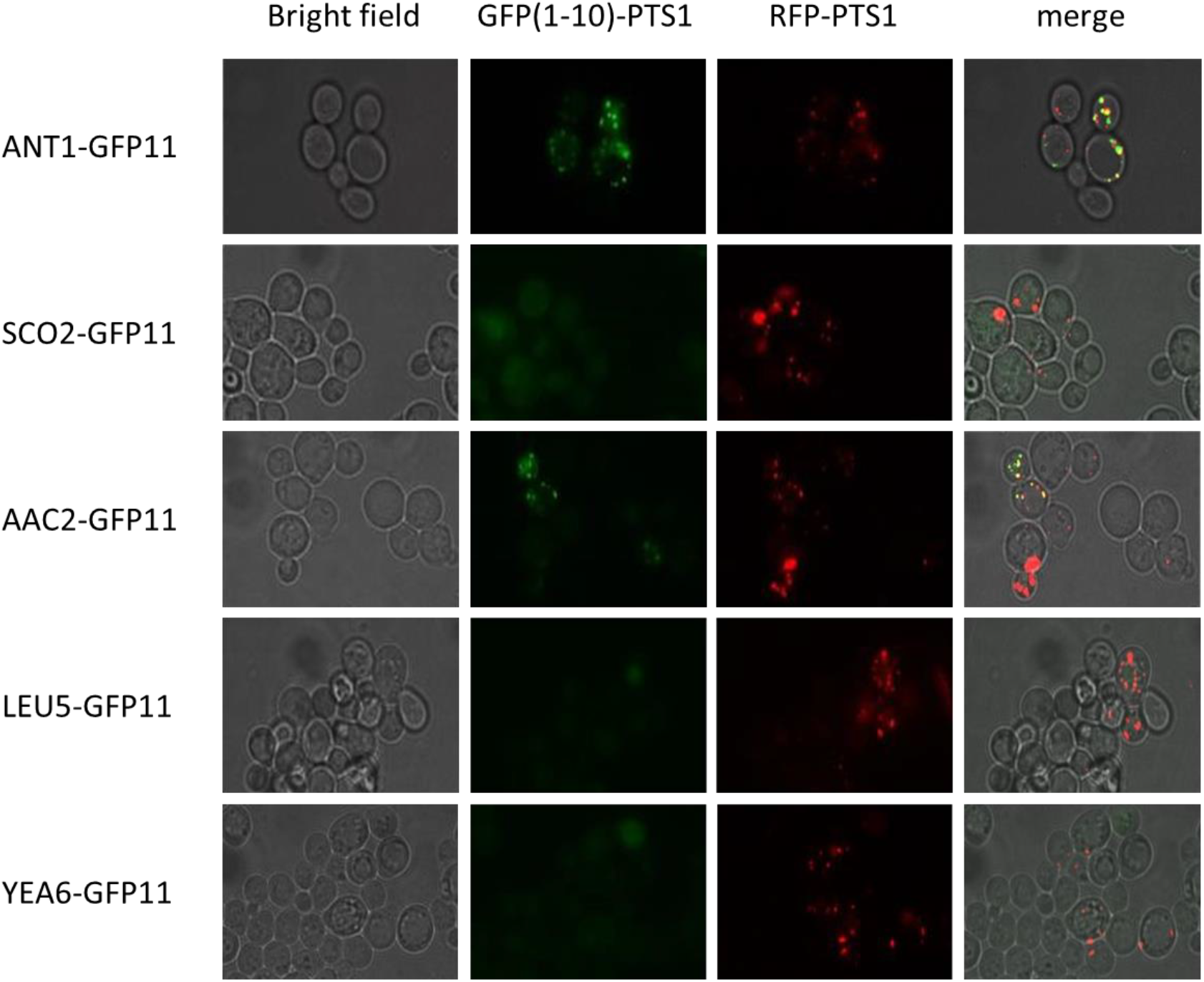
Localisation of MCF proteins using self-assembling GFP assay. Wild-type cells were transformed with GFP(1-10)-PTS1, GFP(11)-tagged MCF proteins (Ant1p, Aac2p, Leu5p, Yea6p or Sco2p) as well as RFP-PTS1 for confirmation of peroxisomal localisation. The peroxisomal ATP transporter (Ant1p) was used as positive control for the assay. All cells were cultured overnight in ethanol medium. Images show bright field to visualize the localization of the cells (left); fluorescence of self-assembling GFP (GFP(1-10)-PTS1; left centre), RFP-PTS1 (right centre) and the overlay of self-assembling GFP and RFP-PTS1 (Merge; right).

### Increased mitochondrial targeting of Aac2p diminishes residual peroxisomal ATP levels

In order to demonstrate that peroxisome-localized Aac2p is responsible for the residual C8:0 β-oxidation activity in the *ant1*Δ *pxa1*Δ *pxa2*Δ mutant cells, we attempted to delete the *AAC2* gene in these cells, but this did not result in a viable strain. As an alternative approach, we introduced a strong mitochondrial targeting signal (MTS) to the N terminus of Aac2p41 in order to increase mitochondrial and decrease peroxisomal targeting of Aac2p. Using our self-assembling GFP assay, we observed that the addition of the strong MTS indeed reduced the peroxisomal localization of GFP(11)-tagged Aac2p to 2-3% of cells compared to 15% of GFP(11)-tagged Aac2p without the MTS (Fig. 5).

**Figure 5.**
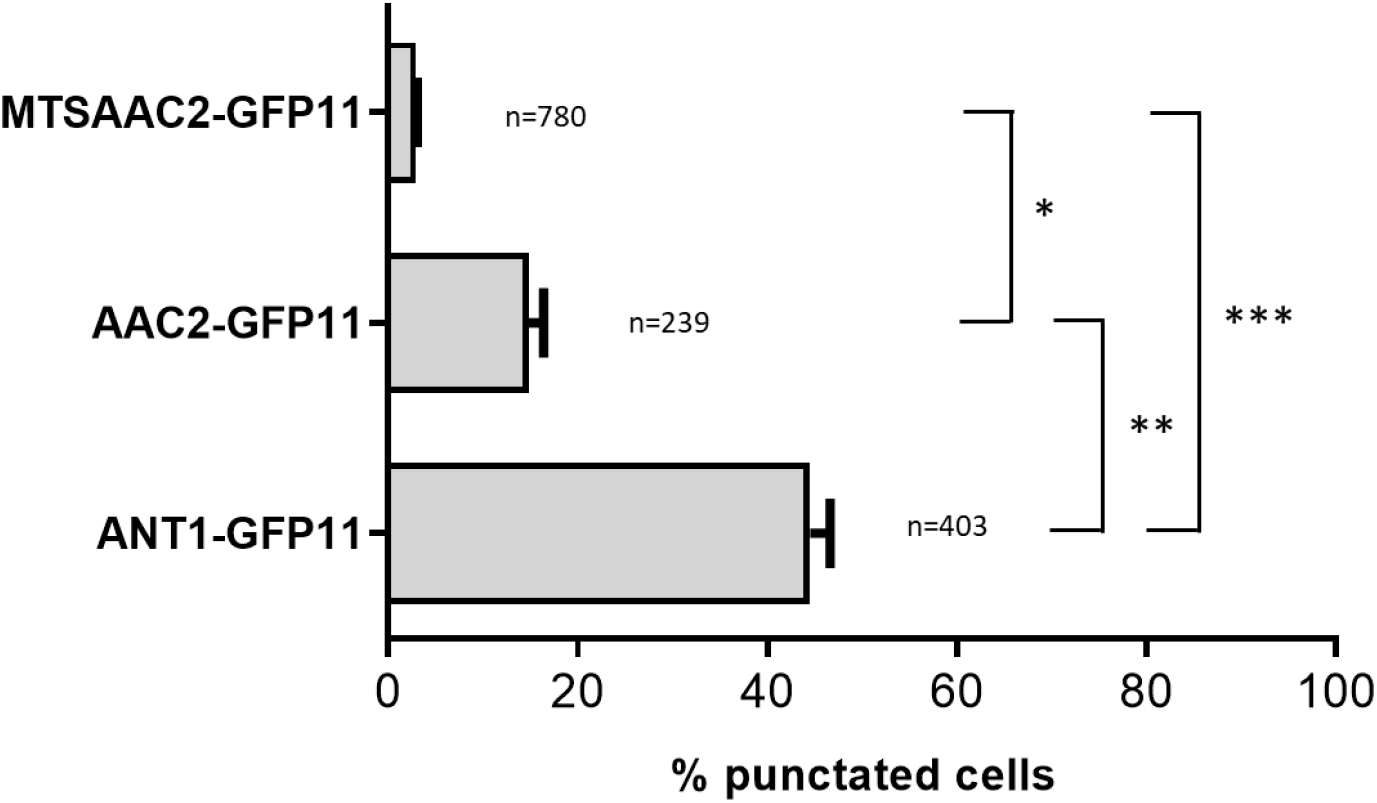
Quantification of cells with Ant1p, Aac2p or MTSAac2p located in the peroxisomal membrane. Wild-type cells co-expressing the indicated GFP(11)-tagged MCF protein and the peroxisomal GFP(1-10)-PTS1 protein were analyzed for the number of cells displaying punctated GFP fluorescence, resulting from the physical interaction between GFP(11)-tagged MCF proteins and the peroxisomal GFP(1-10)-PTS1 protein. Expressed as percentage of number of analyzed cells indicated next to the bars. A one-way ANOVA test with Tukey’s multiple comparisons test was performed for n=3. ***, ** and * indicate significance with a P-value of p<0.001, p<0.01 and p<0.05 respectively.

Next, we introduced the MTS sequence via homologous genomic recombination to the N terminus of Aac2p in wild-type cells, and the *ant1Δ* and *ant1Δ pxa1Δ pxa2Δ* mutant cells. In all three strains, the expression of MTSAac2p resulted in markedly lower C8:0 β-oxidation activities when compared to the activities in the same strains that do not express MTSAac2p (Fig. 6). These findings strongly suggest that peroxisome-localized Aac2p is responsible for the residual C8:0 β-oxidation activity observed in the *ant1Δ pxa1Δ pxa2Δ* mutant cells. When we measured the relative ATP levels using the ATeam reporter proteins, however, we did not observe a significant difference between the peroxisomal ATP levels in wild-type and *ant1Δ* cells when compared to the same cells expressing MTSAac2p (Fig. 6B). This could suggest that the activity of Pxa1/Pxa2p may be sufficient to maintain the ATP levels in *ant1Δ* cells expressing MTSAac2p, but it cannot be excluded that the ATeam reporter proteins are not sensitive enough to detect potential small differences in ATP levels between wild-type or *ant1Δ* cells with and without MTSAac2p. The effect of MTSAac2p on the peroxisomal ATP levels in *ant1Δ pxa1Δ pxa2Δ* mutant cells could not be studied since the residual ATP levels in this mutant strain were below the detection limit of the ATeam reporter proteins.

**Figure 6.**
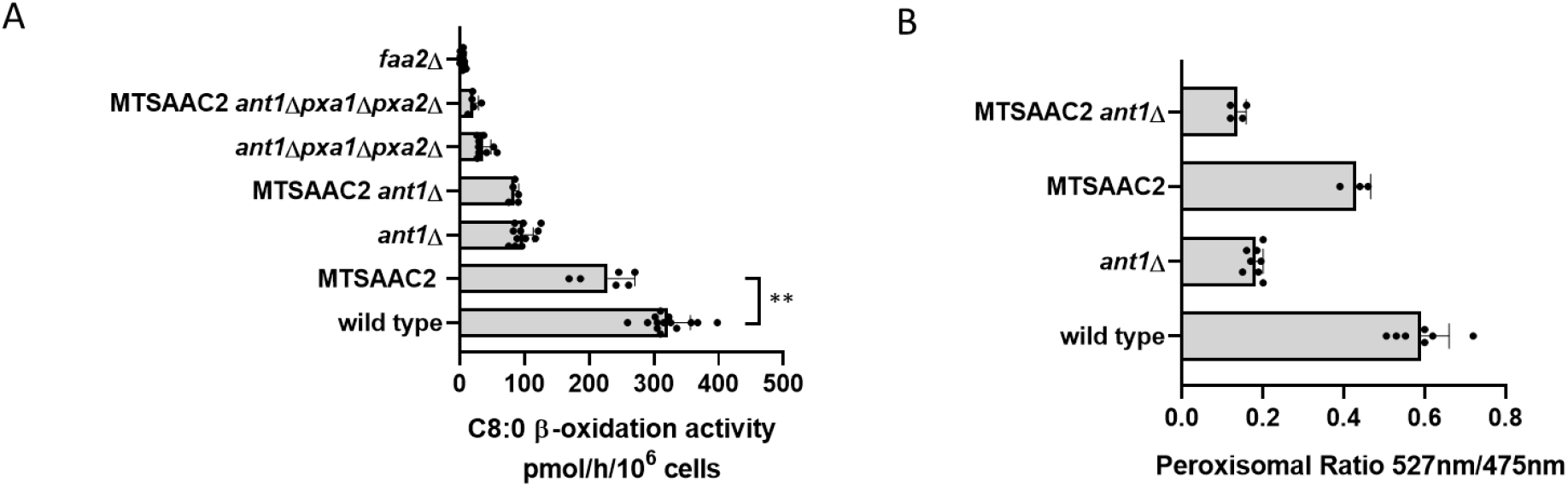
C8:0 β-oxidation activity and ATP levels in wild-type and mutant yeast strains expressing MTS-Aac2p. (**A**) Untransformed strains and the same strains expressing MTSAac2p were cultured overnight in oleate medium and β-oxidation rates were measured using [1-^14^C] labelled octanoate (C8:0) as substrate. Data are means ± SD of values from 5–14 independent experiments. (**B**) Untransformed wild-type and *ant1Δ* strains and the same strains expressing MTSAac2p were transformed with ATeam-per or ATeam-permut to determine relative peroxisomal ATP levels as described in legend of figure 3. Data are means ± SD of values from 3–8 independent experiments. A one-way ANOVA test with Tukey’s multiple comparisons test was performed. ** indicate significance with a P-value of p<0.01.

### Human ABCD1, ABCD2 and ABCD3 can also transport ATP

After having established that Pxa1p/Pxa2p can mediate ATP uptake into peroxisomes, we studied whether the human orthologues HsABCD1, HsABCD2 and HsABCD3 can also mediate peroxisomal ATP uptake in addition to transport of fatty acyl-CoAs. We previously showed that yeast-codon optimized HsABCD1, HsABCD2 and HsABCD3 can be functionally expressed as homodimers in *S. cerevisiae*, and display different substrate specificities^42^.

Expression of HsABCD1 in *ant1Δ pxa1Δ pxa2Δ* mutant cells resulted in a more than 3-fold increase in C8:0 β-oxidation activity (Fig. 7A) and co-expression with the peroxisomal ATeam reporter proteins showed a marked increase in peroxisomal ATP levels (Fig. 7D). Thus, similar as its yeast orthologues, HsABCD1 can also mediate ATP uptake. Expression of ABCD2 or ABCD3 in *ant1Δ pxa1Δ pxa2Δ* mutant cells did not result in a significant increase in C8:0 β-oxidation activity (Fig. 7A).

**Figure 7.**
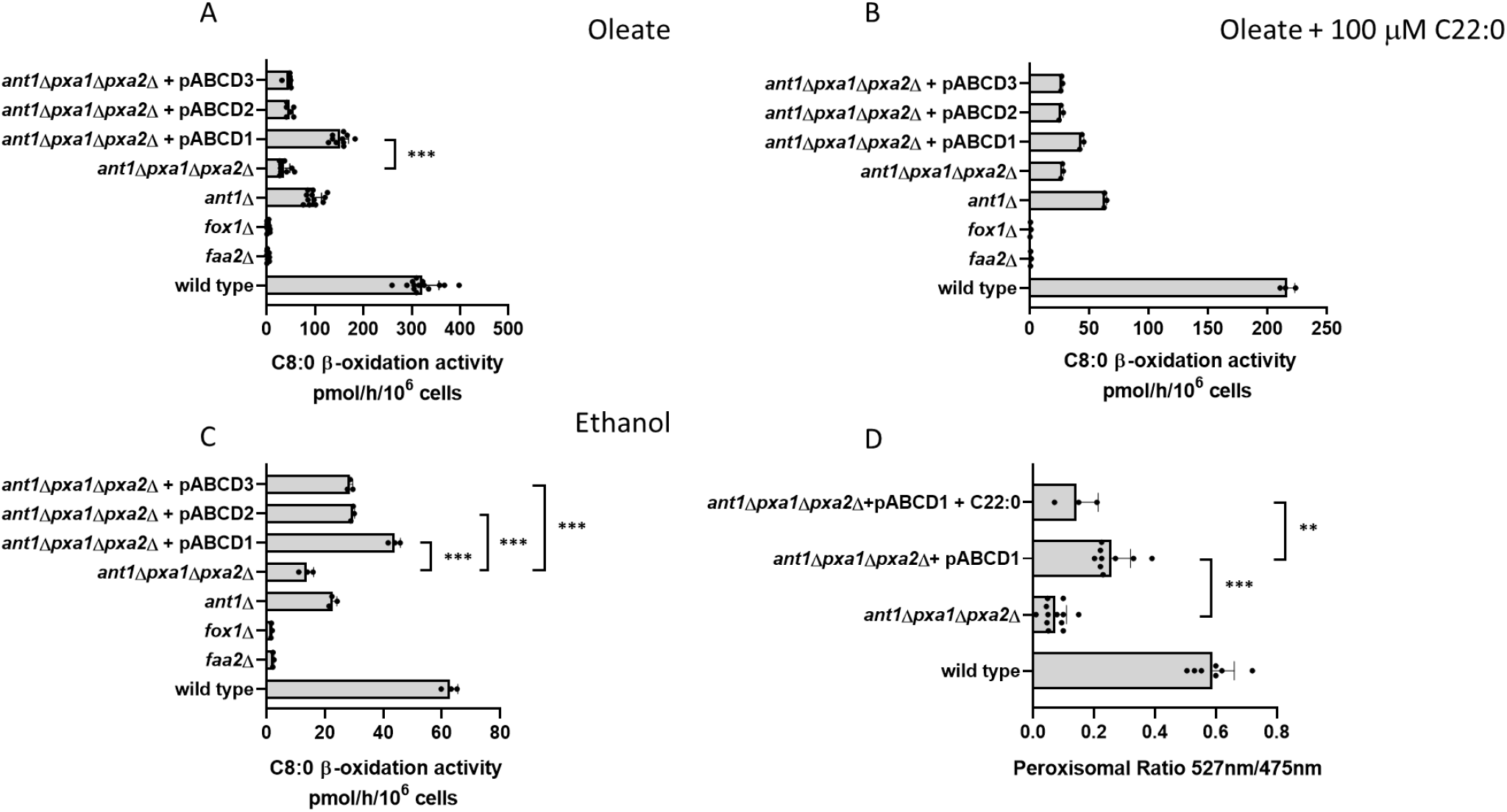
C8:0 β-oxidation activity in wild-type and mutant yeast strains expressing human ABCD proteins. Wild-type and mutant strains, including *ant1Δpxa1Δpxa2Δ* strains expressing ABCD1, ABCD2 or ABCD3 were cultured overnight in oleate medium (**A**), oleate medium supplemented with 100 μM C22:0 (**B**), or ethanol medium (**C**). Fatty acid ß-oxidation activity was measured with [1-^14^C] labelled octanoate (C8:0) (**A, B, C**). (**D**) Wild-type and mutant strains, including *ant1Δpxa1Δpxa2Δ* strains expressing ABCD1 were transformed with ATeam-per and ATeam-permut to allow quantification of peroxisomal ATP levels. Cells were grown overnight in oleate medium or oleate medium supplemented with 100 μM C22:0. ATP levels were determined as described in legend of Figure 3. Data are means ± SD of values from 3–12 independent experiments. A one-way ANOVA test with Tukey’s multiple comparisons test was performed. ***, ** and * indicate significance with a P-value of respectively p<0.001, p<0.01 and p<0.05.

The substrate preference of ABCD1 for very-long-chain acyl-CoAs allowed us to study in yeast whether ATP and acyl-CoA compete for the same binding site of ABCD1. To this end, we measured C8:0 β-oxidation activity in the presence and absence of docosanoic acid (C22:0), which is a good substrate for ABCD1 after intracellular conversion into its CoA-ester^42^. Both the C8:0 β-oxidation activity (Fig. 7B) and the peroxisomal ATP levels (Fig. 7D) were lower in the presence of C22:0, which strongly suggests that the fatty acyl-CoAs and ATP compete for the same binding site in HsABCD1.

All experiments described so far were performed with yeast cells cultured on oleate (C18:1), which is also substrate for the ABCD proteins and thus could compete for the ATP uptake by the ABCD proteins similar as C22:0. To prevent such competition, we cultured the *ant1Δ pxa1Δ pxa2Δ* mutant cells expressing HsABCD1, HsABCD2 or HsABCD3 on ethanol instead of oleate and then repeated the C8:0 β-oxidation activity measurements. We now measured a significant increase in C8:0 beta-oxidation activity in the cells expressing HsABCD1 as well as in the cells expressing HsABCD2 or HsABCD3 when compared to the same cells expressing neither of these proteins (Fig. 7C). Taken together, these results show that all three human ABCD transporters can mediate peroxisomal ATP uptake in addition to transport of fatty acyl-CoAs.

## Discussion

Peroxisomes are generally considered to be selectively permeable organelles, which allow molecules with an Mw<400Da to cross the peroxisomal membrane passively via one or more (putative) channel-forming proteins, including Pxmp2^20^ and Pex11beta^29,30^. Transport across the peroxisomal membrane of more bulky molecules, such as fatty acyl-CoAs and ATP, however, requires dedicated transport proteins. Among others, intra-peroxisomal ATP is essential for the peroxisomal β-oxidation of fatty acids. Previous work has shown that the transport of fatty acyl-CoAs is mediated by different dimeric half-ABC transporters of the ABCD family, which includes the heterodimer Pxa1p/Pxa2p in peroxisomes of *S. cerevisiae* and homodimers of ABCD1, ABCD2, and ABCD3 in human peroxisomes. In this paper we have studied which proteins are involved in the peroxisomal uptake of ATP. ATP has to be transported from the cytosol into the peroxisome since peroxisomes do not possess an ATP-synthesizing or regenerating system. To this end, we generated and used a series of *S. cerevisiae* mutant strains in which we deleted single or combinations of genes encoding putative peroxisomal ATP transporters. As readouts for peroxisomal ATP uptake we used three different and independent assays, including i) measurement of C8:0 β-oxidation activity, which in yeast is strictly dependent on intra-peroxisomal ATP; ii) ATP uptake/exchange by proteoliposomes prepared from peroxisomal membranes isolated from the different mutant strains; and iii) measurement of relative ATP levels using FRET-based ATeam reporter proteins targeted to the peroxisomes or the cytosol.

We demonstrated that peroxisomes in *S. cerevisiae* contain three different transport systems that can mediate ATP uptake, each with a different mode of action. As anticipated, we found the previously reported peroxisomal membrane protein Ant1p to be responsible for most of the peroxisomal ATP uptake as can be concluded from the observation that a deletion of *ANT1* resulted in a ~70% reduction of the peroxisomal ATP-dependent C8:0 β-oxidation activity and in a marked reduction of the intra-peroxisomal ATP levels. As reported previously and confirmed in our proteoliposome uptake studies, Ant1p most probably functions as an ATP/AMP antiporter^27, 28^. Surprisingly, we found that the remaining peroxisomal ATP uptake capacity is mediated by the ABC transporter protein complex Pxa1p/Pxa2p, which most probably functions as an ATP uniporter, and Aac2p, a predominantly mitochondrial ATP/ADP antiporter^33,38,39,40^, which we here showed to be partly localized in peroxisomes.

Our finding that heterodimeric Pxa1p/Pxa2p, as well as the human orthologues HsABCD1, HsABCD2 and HsABCD3 as homodimers, can also mediate peroxisomal uptake of non-hydrolyzed ATP in addition to fatty acyl-CoAs was unexpected and is very intriguing. Indeed, ABC transporter proteins typically use ATP hydrolysis to catalyze the transport of substrates across membranes, although few ABC transporter proteins have been reported to function as ATP channels^43, 44, 45, 46^. Our substrate competition experiments in which we added C22:0 during the C8:0 β-oxidation measurement indicated that ATP and acyl-CoAs compete for the same binding site of HsABCD1. This most likely is different from the binding site at which ATP is hydrolyzed to drive the transport of fatty acyl-CoAs across the peroxisomal membrane.

Aac2p is a well-established ATP/ADP antiporter localized in the inner mitochondrial membrane^33,38–40^. Our finding that Aac2p is partly localized in peroxisomes and thus constitutes a third transport protein mediating peroxisomal ATP uptake is equally intriguing as the finding of the involvement of Pxa1p/Pxa2p. It also raises the question on how Aac2p is targeted to peroxisomes in addition to its primary targeting to mitochondria, given that the mechanisms of membrane protein import into mitochondria and peroxisomes must be very different, although this is still largely unknown for peroxisomal membrane proteins. It should be noted, however, that a dual localization in mitochondria and peroxisomes is not unique and has been reported for several proteins, including DLP1, FIS1, MIRO, and VDAC^47^. In the case of Aac2p, the introduction of a stronger mitochondrial targeting signal (MTS) to its N terminus reduced the peroxisomal localization and, consequently, the peroxisomal ATP uptake mediated by peroxisomal Aac2p.

The different substrate affinities and modes of action of Ant1p, Pxa1p/Pxa2p and Aac2p not only ensure that peroxisomes can maintain their ATP levels to support the intra-peroxisomal ATP-dependent enzyme reactions, but also take care of the export of AMP and ADP generated after intra-peroxisomal hydrolysis of ATP. Indeed, peroxisomes in yeast harbor several enzymes the activity of which depends on ATP hydrolysis leading to the generation of AMP or ADP. These include several acyl-CoA synthetases^17, 23^, PCD1^48^, NPY1^49^, VPS34 kinase^50^ and LonP proteases^51, 52^.

Taken together, our results provide compelling evidence for the presence of multiple systems for the transport and exchange of ATP in peroxisomes in yeast. Our finding that, similar as their yeast orthologs Pxa1p/Pxa2p, also the human peroxisomal ABC transporters can mediate peroxisomal ATP uptake, strongly suggests that these findings are transferable to human peroxisomes.

## Methods

### Yeast strains

We used *S. cerevisiae* BJ1991 (*MATα, pep4-3, prbl-1122, ura3-52, leu2, trp1*) as wild-type strain and for the generation of targeted deletion mutant strains. Gene deletions in BJ1991 were created by replacement of specific genes by the yeast *LEU2* gene, the Kanamycin (*KAN*) or the Bleomycin (*BLE*) resistance gene using homologous recombination. For this study we generated and used the following deletion mutant strains: *ant1Δ* (*ant1*::*KAN*), *pxa1Δ pxa2Δ* (*pxa1*::*LEU2*, *pxa2*::*KAN*), and *ant1Δ pxa1Δ pxa2Δ* (*ant1*::*KAN*, *pxa1*::*LEU2*, *pxa2*::*BLE*), and used two previously described mutant strains *faa2Δ* (*faa2*::*LEU2*) and *fox1Δ* (*fox1*::*KAN*).

### Culture conditions

We cultured yeast cells at 28°C under continuous shaking at 225 rpm. For standard growth, cells were cultured in glucose medium containing 6.7 g/L yeast nitrogen base without amino acids (Difco) and 5 g/L D-glucose. Amino acids were supplemented to the medium when required; 30 mg/L leucine, 20 mg/L uracil, or 20 mg/L tryptophan. To induce peroxisome proliferation, yeast cells were cultured for at least 24 hours in glucose medium and then transferred to and cultured overnight in YPO medium (3 g/L yeast extract, 5 g/L peptone, 25 mM potassium phosphate buffer (pH=6), 1.07 g/L oleate, 2.16 g/L Tween-80) with supplemented amino acids when required.

For fluorescent microscopy of self-assembling GFP, yeast cells were cultured in ethanol medium containing 1 g/L yeast extract, 25 mM potassium phosphate buffer (pH=6), 6.7 g/L yeast nitrogen base and 20 ml/L ethanol and supplemented amino acids when required.

### Octanoate (C8:0) β-oxidation measurements

We measured β-oxidation activity in intact yeast cells as follows. Cells were cultured overnight in YPO media, harvested by centrifugation, washed and resuspended in 9 g/L NaCl at a cell density of OD_600nm_=1 (~1.48 × 10^7^ cells). Incubations were performed in 20 mL vials with a rubber septum, containing two tubes, one with the cells in incubation mixture and the other with 500 μL NaOH (2M). To start the measurements, 20 μL of cell suspension was added to the reaction mixture composed of 20 μL MES buffer (0.5M; pH=6), 140 μL NaCl (9 g/L), and 20 μL of 100 μM [1-^14^C] octanoate (200,000 dpm) as substrate. The reaction was allowed to proceed for 1 hour at 28°C after which the reaction was terminated by the addition of 50 μL of perchloric acid (2.6 M). Radiolabelled [^14^C]-CO2, released during the β-oxidation of octanoate and trapped in the tube with NaOH, was counted in a liquid scintillation counter and used to quantify the rate of β-oxidation in nmol/h/107 cells. The rate of octanoate β-oxidation in wild-type cells was 3.2 ± 0.4 nmol/h/10^7^ cells.

### ATP uptake measurements in proteoliposomes

We isolated peroxisomes in duplicate from wild-type and the *ant1Δ*, *pxa1Δpxa2Δ* and *ant1Δpxa1Δpxa2Δ* mutant strains cultured overnight in oleate medium using cell fractionation and Nycodenz gradient centrifugation as described previously^28^. Gradient fractions were analysed for peroxisomal 3-hydroxyacyl-CoA dehydrogenase (3-HAD) and mitochondrial fumarase activity^28^. Purified peroxisomes from fractions 2-4 of the gradients (Fig. S1) and equivalent to 375 units of peroxisomal 3HAD activity were harvested and dissolved in 150 μL of 50mM Hepes (pH=7.4) and 5 mM MgCl_2_. Of these, peroxisomes equivalent to 50 units of 3HAD activity were added to 1 mL 30 g/L L-α-phosphatidylcholine only or supplemented with 10 mM ADP or 10 mM AMP after which the mixtures were frozen in liquid nitrogen. The samples were then thawed at room temperature, resulting in the formation of proteoliposomes, and subjected to size-exclusion chromatography using Sephadex G-25 (Medium) columns (GE Healthcare Life Science) to remove external ADP or AMP. The eluate was used to start the uptake experiment by adding 0.2 mM [α-^32^P]-ATP (6,000 Ci/mmol). The uptake reaction was terminated via passing the proteoliposomes over Dowex AG1-X8 anion-exchange columns using 150 mM sodium acetate (pH7.4) as elution buffer. The incorporated [α-^32^P]-ATP was quantified by liquid scintillation counting. Time-dependent uptake data were fitted using nonlinear regression analysis based on one-phase exponential association using GraphPad Prism 5.0 software (GraphPad, www.graphpad.com). The initial velocity of uptakes were calculated using the equation slope = (Plateau - Y0)*k, with Y0 set to 0.

### Measurement of ATP levels using FRET-based ATeam reporter proteins

We measured the relative *in vivo* ATP levels in peroxisomes and the cytosol of wild-type cells and different mutant strains through expression of modified versions of the previously described ATeam sensors^35^. As source for the generation of the ATeam reporter proteins used in this study, we ordered the pDR-GW AT1.03 and pDR-GW AT1.03 R122K/R126K plasmids^36^ from Addgene (deposited by Wolf Frommer). The pDR-GW AT1.03 plasmid (http://www.addgene.org/28003) codes for a cytosolic ATeam reporter protein and the pDR-GW AT1.03 R122K/R126K plasmid (http://www.addgene.org/28005) codes for a mutated version of the same ATeam reporter protein that no longer binds ATP. To allow constitutive, carbon source-independent Ateam gene expression in yeast, we first replaced the CTA1 promoter of pIJL30 with the TEF1 promoter generating the yeast expression vector pMK05 (TEF1pr, ARS1/CEN4, Trp1, ampR). The *XbaI*-*HindIII* fragments from pDR-GW AT1.03 and pDR-GW AT1.03R122K/R126K were then subcloned downstream of the TEF1 promoter into the *XbaI*-*HindIII* sites of pMK05. The resulting plasmids were designated pATeamcyt (expressing cytosolic AT1.03) and pATeamcytmut (expressing the mutated cytosolic AT1.03 R122K/R126K), respectively.

To target the ATeam reporter proteins to peroxisomes, we replaced by site directed mutagenesis the stop codon of the AT1.03 ORFs in pATeamcyt and pATeamcytmut by a flexible loop and the coding sequence for the twelve C-terminal amino acids of Fox2p, including the strong peroxisomal targeting sequence PTS1. This resulted in pATeamper (expressing peroxisomal AT1.03) and pATeampermut (expressing the mutated peroxisomal AT1.03 R122K/R126K), respectively.

Yeast cells were transformed with the different pATeam plasmids and cultured overnight in YPO medium. The cells were harvested by centrifugation at 230 *g* for 5 minutes at 4°C, washed once with and then suspended in cold MES-glucose buffer (pH 6.0) composed of 25 mM 2-(N-morpholino) ethanesulfonic acid and 20 g/L D-glucose. Cells were kept on ice until analysis was conducted (5 hours max). Prior to the measurements, the suspended cells were diluted with MES-glucose buffer until OD_600nm_=3, and 200 μL was transferred in duplicate into a 96-wells microplate (Grenier, black round bottom). FRET analysis was then performed in on a Tecan Infinite M200 pro plate reader using an excitation of 435/9 nm and detecting emission at 475/20 nm and 527/20 nm, respectively. Fluorescence intensities at each wavelength were measured 10 times. To assure accurate measurements, prior to the start of each cycle the microplate was shaken in orbitals with an amplitude of 2 mm, a Z position height of 20000 μm, settle time of 200 ms, and 0 s lag time. Ratios were calculated after substraction of the background signal in wild-type cells transformed with an empty plasmid (no Ateam expression). The relative ATP levels were obtained by subtracting the 572/475 ratio of the Ateamcyt/per (mut) reporter protein from the 572/475 ratio of the corresponding Ateamcyt/per reporter protein.

### Construction of ABCD expression plasmids

We designed and ordered a yeast-codon optimized open reading frame (ORF) coding for ABCD1 and flanked by *SacI* and *KpnI*, and cloned this into the yeast expression vectors pIJl30 and pEL30. Construction of ABCD2 and ABCD3 expression plasmids have been described previously^42^.

### Subcellular localisation of MCF proteins using a self-assembling GFP assay

We adapted the self-assembling GFP assay^37^ to study the subcellular localization of MCF proteins. To this end, we designed and ordered from Genscript a yeast-codon optimized open reading frame (ORF) coding for GFP(1-10) in pUC57 and re-cloned this into the yeast expression vector pIJl30 (CTA1pr, ARS1/CEN4, Trp1, ampR) allowing cytosolic expression of GFP(1-10). To generate a peroxisome-localized GFP(1-10), we used the pUC57-GFP(1-10) as template and added via PCR amplification the coding sequences of the twelve C-terminal amino acids of FOX2, which include a strong peroxisomal targeting signal PTS1, spaced with a flexible linker. The resulting GFP(1-10)-PTS1 ORF was also subcloned into the pIJl30 expression vector. To generate a basic cloning vector that allows expression of MCF proteins with a C-terminal extension coding for the GFP(11) domain, we introduced a linker encoding a flexible loop and yeast-codon optimized GFP(11) into the *BamH1* and *HindIII* sites of yeast expression vector pEL3017 (CTA1pr, ARS1/CEN4, URA3, ampR). The resulting plasmid pEL30-GFP(11) allows upstream cloning of ORFs into *SacI*, *KpnI*, *SmaI* and *BamHI* sites in frame with GFP(11).

ORFs encoding the yeast MCF proteins Ant1p, Aac2p, Yea6p, Leu5p and Sco2p were PCR amplified from genomic DNA of *S. cerevisiae* using ORF-specific PCR primers with small extensions to introduce the appropriate restriction sites and, after restriction, sub-cloned in frame with GFP(11) into pEL30-GFP(11). All PCR-amplified sequences were verified by Sanger sequencing.

Wild-type cells were co-transformed with pEL30-GFP(11) containing one of the MCF proteins and either pIJL30-GFP(1-10)-PTS1 or pIJL30-GFP(1-10). After transformation the cells were cultured in 2% ethanol medium for 24 hours, harvested, re-suspended in sterile water and examined on a ZEISS Axio Observer A1 fluorescence microscope using a 450 nm excitation and a 515-565 nm emission filter. The Leica Application Suite was used to capture the images.

### Enhancing mitochondrial targeting of Aac2p

We introduced the mitochondrial targeting signal (MTS) from the mitochondrial succinate /fumarate carrier of *Arabidopsis thaliana* to the N-terminus of Aac2p^41^, to increase mitochondrial and decrease peroxisomal targeting of Aac2p. To this end, we amplified the ORF of *AAC2* by PCR using an *AAC2*-specific forward primer with a 5’ extension comprising the coding sequence for the MTS. The MTS-*AAC2* ORF was cloned in frame with GFP(11) into pEL30-GFP(11) and verified by Sanger sequencing. Wild-type cells were co-transformed with the resulting pEL30-MTS-Aac2p-GFP(11) vector and the pIJL30-GFP(1-10)-PTS1 vector and used in the self-assembling GFP assay described above to compare the subcellular localization of MTS-Aac2p with Aac2p.

In order to introduce the MTS at the N terminus of genomically encoded Aac2p by homologous recombination, we generated by PCR amplification a DNA fragment comprising a 5’ *AAC2* non-coding sequence followed by the *NAT1* resistance gene under control of the *TEF1* promoter, the sequence for the MTS under control of the *NOP1* promoter, and a 5’ *AAC2* coding sequence. The fragment was transformed into wild-type, *ant1Δ* and *ant1/pxa1/pxa2Δ* mutant strains. After transformation, cells were washed and incubated for 5 hours in 5 g/L glucose supplemented with amino acids, so that the *NAT1* resistance gene could be expressed. Cells were then plated on YPD plates supplemented with 100 μg/mL NTC to select for cells expressing the *NAT1* gene. Correct integration of the DNA fragment at the *AAC2* locus was verified by Sanger sequencing. Normal growth was observed in all knock-in strains on either YPD or 5 g/L glucose medium supplemented with amino acids. The different knock-in strains were used to determine the relative peroxisomal and cytosolic ATP levels using the ATeam reporter constructs as described above.

## Acknowledgements

We thank Suzan Knottnerus, Andrew McDonald, Juan Carlos Munoz, Demi Rijlaarsdam and Mark de Keizer for their input in parts of this study. This work was in part funded by the German Research Foundation (DFG) via grants LI 1781/1-3 (to NL).

## Author contributions

C.v.R and L.IJ. designed research and analysed data.

C.v.R. and N.L. performed experiments.

C.v.R, L.IJ, R.W., N.L. and H.W. wrote and revised the manuscript.

All authors declare no competing interest.

---

**Figure S1.**
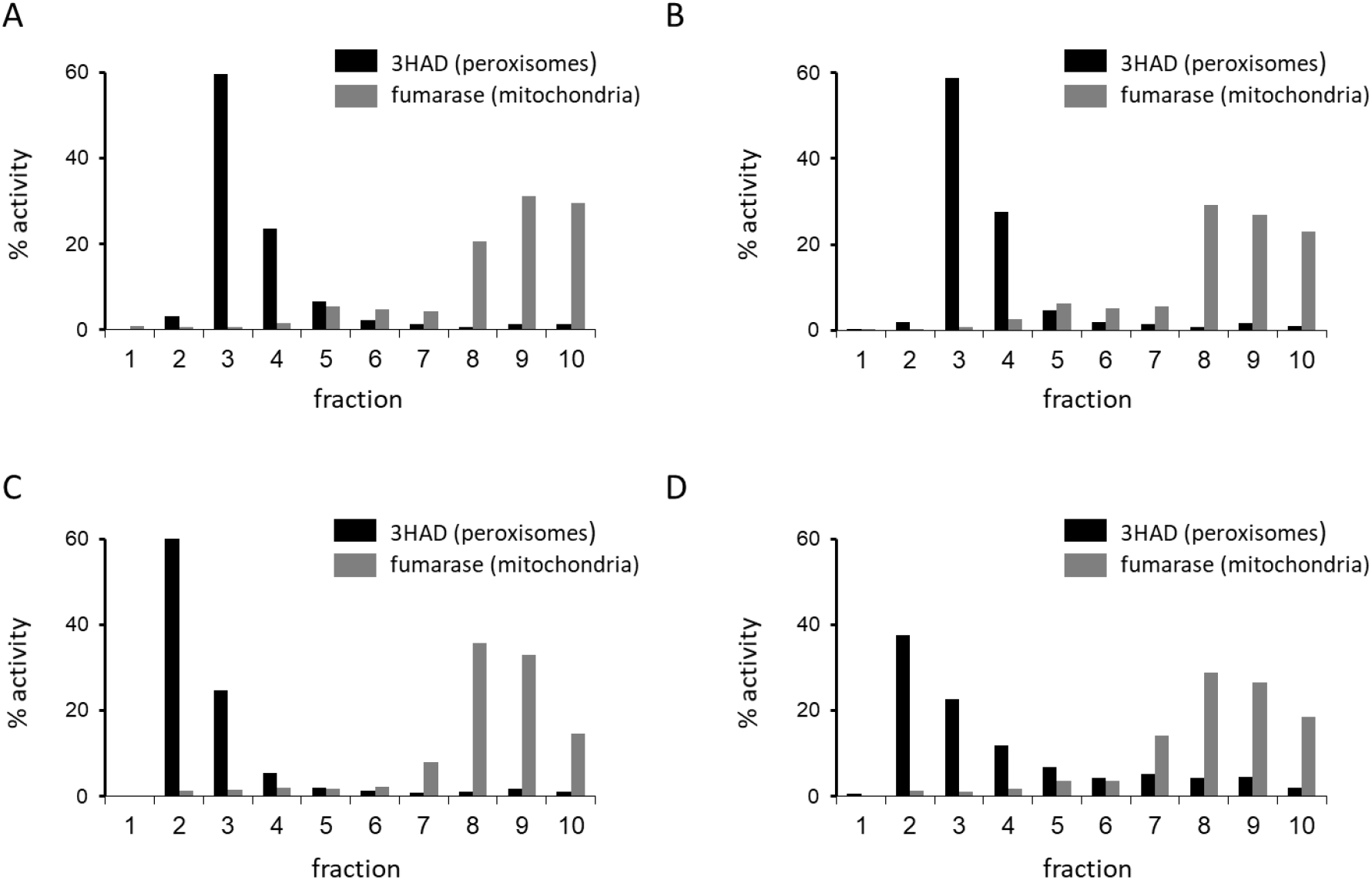
Isolation of peroxisomes from oleate grown yeast strains used for preparation of proteoliposomes. Wild-type (**A**), *ant1Δ* (**B**), *pxa1Δpxa2Δ* (**C**) and *ant1Δpxa1Δpxa2Δ* (**D**) strains were cultured overnight in oleate rich medium and peroxisomes were isolated after cell fractionation and nycodenz gradient centrifugation. A 100.000 *g* pellet of fraction 2, 3 and 4 was used to prepare proteoliposome for *in vitro* transport assays as shown in figure 2.

**Figure S2.**
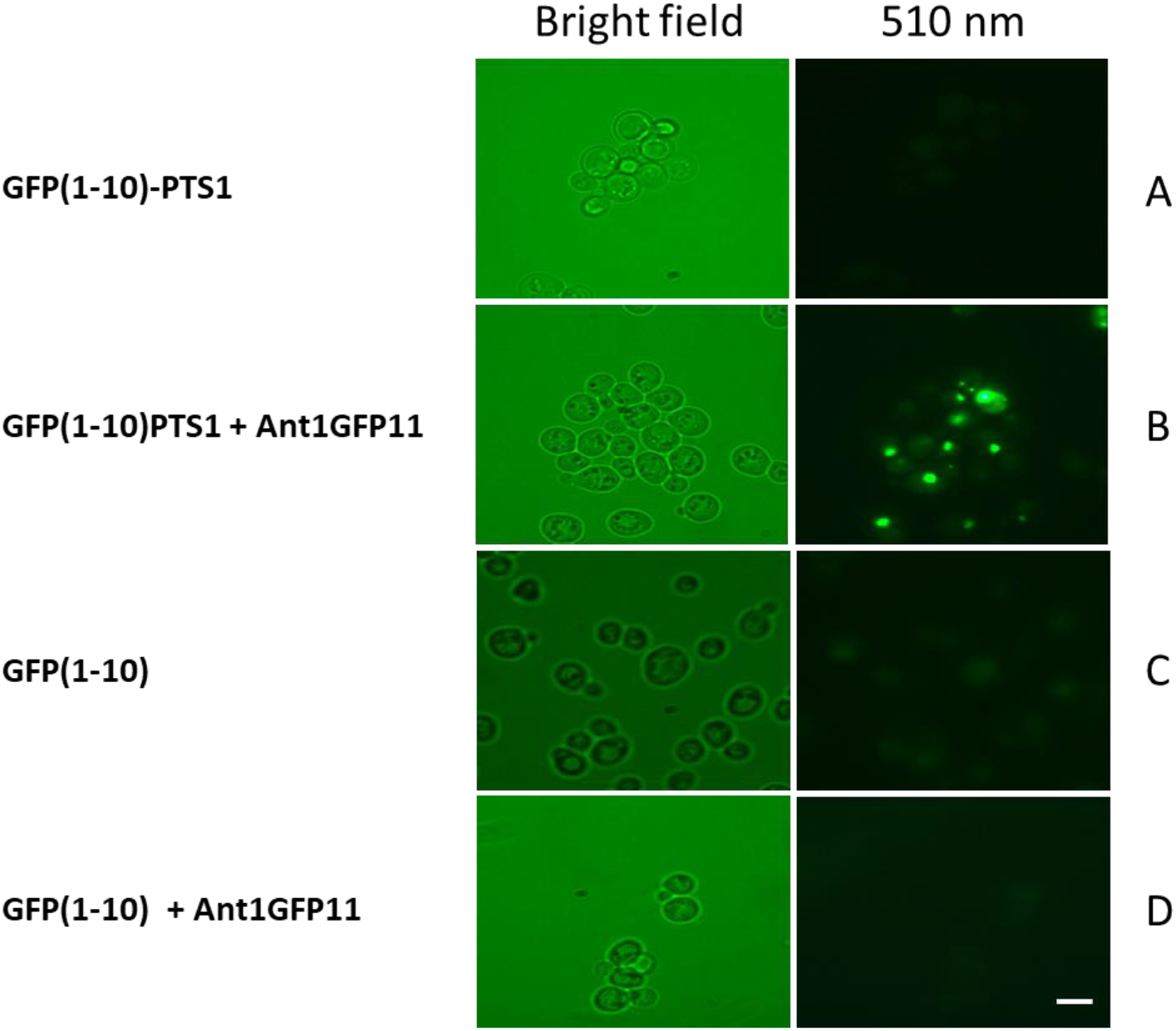
Validation of self-assembling GFP assay. To validate the self-assembling GFP assay used for localizing MCF proteins to peroxisomes, we expressed a peroxisome-targeted GFP(1-10)-PTS1 (panel A+B) and a cytosol-targeted GFP(1-10) protein in wild-type cells, without (panel A and C) and with GFP(11)-tagged Ant1p protein (panel B and D). GFP fluorescence due to physical interaction between GFP(11) and GFP(1-10) only occurred in cells expressing peroxisomal GFP(1-10)-PTS1 and Ant1-GFP(11) (panel B). Scale bar represent 4 μm.

## References

1. Wanders RJ, Waterham HR. Biochemistry of mammalian peroxisomes revisited. Annual review of biochemistry 75, 295–332 (2006).

2. Van Veldhoven PP. Biochemistry and genetics of inherited disorders of peroxisomal fatty acid metabolism. Journal of lipid research 51, 2863–2895 (2010).

3. Waterham HR, Ferdinandusse S, Wanders RJ. Human disorders of peroxisome metabolism and biogenesis. Biochimica et biophysica acta 1863, 922–933 (2016).

4. Wanders RJA. Peroxisomal disorders: Improved laboratory diagnosis, new defects and the complicated route to treatment. Molecular and cellular probes, (2018).

5. Wanders RJ. Metabolic functions of peroxisomes in health and disease. Biochimie 98, 36–44 (2014).

6. Mi J, Kirchner E, Cristobal S. Quantitative proteomic comparison of mouse peroxisomes from liver and kidney. Proteomics 7, 1916–1928 (2007).

7. Wiese S, et al. Proteomics characterization of mouse kidney peroxisomes by tandem mass spectrometry and protein correlation profiling. Molecular & cellular proteomics : MCP 6, 2045–2057 (2007).

8. Gronemeyer T, et al. The proteome of human liver peroxisomes: identification of five new peroxisomal constituents by a label-free quantitative proteomics survey. PloS one 8, e57395 (2013).

9. Plett A, Charton L, Linka N. Peroxisomal Cofactor Transport. Biomolecules 10, (2020).

10. Yi EC, et al. Approaching complete peroxisome characterization by gas-phase fractionation. Electrophoresis 23, 3205–3216 (2002).

11. Chen X, Williams C. Fungal Peroxisomes Proteomics. Sub-cellular biochemistry 89, 67–83 (2018).

12. Aubourg P, Mosser J, Douar AM, Sarde CO, Lopez J, Mandel JL. Adrenoleukodystrophy gene: unexpected homology to a protein involved in peroxisome biogenesis. Biochimie 75, 293–302 (1993).

13. Mosser J, et al. Putative X-linked adrenoleukodystrophy gene shares unexpected homology with ABC transporters. Nature 361, 726–730 (1993).

14. Holzinger A, Kammerer S, Berger J, Roscher AA. cDNA cloning and mRNA expression of the human adrenoleukodystrophy related protein (ALDRP), a peroxisomal ABC transporter. Biochemical and biophysical research communications 239, 261–264 (1997).

15. Lombard-Platet G, Savary S, Sarde CO, Mandel JL, Chimini G. A close relative of the adrenoleukodystrophy (ALD) gene codes for a peroxisomal protein with a specific expression pattern. Proceedings of the National Academy of Sciences of the United States of America 93, 1265–1269 (1996).

16. Kamijo K, Osumi T, Hashimoto T. [PMP70, the 70-kDa peroxisomal membrane protein: a member of the ATP-binding cassette transporters]. Nihon rinsho Japanese journal of clinical medicine 51, 2343–2352 (1993).

17. van Roermund CW, et al. Peroxisomal fatty acid uptake mechanism in Saccharomyces cerevisiae. The Journal of biological chemistry 287, 20144–20153 (2012).

18. Okamoto T, et al. Characterization of human ATP-binding cassette protein subfamily D reconstituted into proteoliposomes. Biochemical and biophysical research communications 496, 1122–1127 (2018).

19. Agrimi G, Russo A, Scarcia P, Palmieri F. The human gene SLC25A17 encodes a peroxisomal transporter of coenzyme A, FAD and NAD+. The Biochemical journal 443, 241–247 (2012).

20. Rokka A, et al. Pxmp2 is a channel-forming protein in Mammalian peroxisomal membrane. PloS one 4, e5090 (2009).

21. Reguenga C, Oliveira ME, Gouveia AM, Eckerskorn C, Sá-Miranda C, Azevedo JE. Identification of a 24 kDa intrinsic membrane protein from mammalian peroxisomes. Biochimica et biophysica acta 1445, 337–341 (1999).

22. Visser WF, van Roermund CW, Ijlst L, Waterham HR, Wanders RJ. Metabolite transport across the peroxisomal membrane. The Biochemical journal 401, 365–375 (2007).

23. Hettema EH, et al. The ABC transporter proteins Pat1 and Pat2 are required for import of long-chain fatty acids into peroxisomes of Saccharomyces cerevisiae. The EMBO journal 15, 3813–3822 (1996).

24. Shani N, Sapag A, Watkins PA, Valle D. An S. cerevisiae peroxisomal transporter, orthologous to the human adrenoleukodystrophy protein, appears to be a heterodimer of two half ABC transporters: Pxa1p and Pxa2p. Annals of the New York Academy of Sciences 804, 770–772 (1996).

25. Swartzman EE, Viswanathan MN, Thorner J. The PAL1 gene product is a peroxisomal ATP-binding cassette transporter in the yeast Saccharomyces cerevisiae. The Journal of cell biology 132, 549–563 (1996).

26. van Roermund CW, et al. The human peroxisomal ABC half transporter ALDP functions as a homodimer and accepts acyl-CoA esters. FASEB journal : official publication of the Federation of American Societies for Experimental Biology 22, 4201–4208 (2008).

27. Palmieri L, Rottensteiner H, Girzalsky W, Scarcia P, Palmieri F, Erdmann R. Identification and functional reconstitution of the yeast peroxisomal adenine nucleotide transporter. The EMBO journal 20, 5049–5059 (2001).

28. van Roermund CW, et al. Identification of a peroxisomal ATP carrier required for medium-chain fatty acid beta-oxidation and normal peroxisome proliferation in Saccharomyces cerevisiae. Molecular and cellular biology 21, 4321–4329 (2001).

29. van Roermund CW, Tabak HF, van Den Berg M, Wanders RJ, Hettema EH. Pex11p plays a primary role in medium-chain fatty acid oxidation, a process that affects peroxisome number and size in Saccharomyces cerevisiae. The Journal of cell biology 150, 489–498 (2000).

30. Mindthoff S, et al. Peroxisomal Pex11 is a pore-forming protein homologous to TRPM channels. Biochimica et biophysica acta 1863, 271–283 (2016).

31. van Roermund CW, Waterham HR, Ijlst L, Wanders RJ. Fatty acid metabolism in Saccharomyces cerevisiae. Cellular and molecular life sciences : CMLS 60, 1838–1851 (2003).

32. Carrier DJ, et al. Mutagenesis separates ATPase and thioesterase activities of the peroxisomal ABC transporter, Comatose. 9, 10502 (2019).

33. van Roermund CWT, l IJ, Baker A, Wanders RJA, Theodoulou FL, Waterham HR. The Saccharomyces cerevisiae ABC subfamily D transporter Pxa1/Pxa2p co-imports CoASH into the peroxisome. FEBS Lett, (2020).

34. Palmieri F, et al. Identification of mitochondrial carriers in Saccharomyces cerevisiae by transport assay of reconstituted recombinant proteins. Biochimica et biophysica acta 1757, 1249–1262 (2006).

35. Imamura H, et al. Visualization of ATP levels inside single living cells with fluorescence resonance energy transfer-based genetically encoded indicators. Proceedings of the National Academy of Sciences of the United States of America 106, 15651–15656 (2009).

36. Bermejo C, Haerizadeh F, Takanaga H, Chermak D, Frommer WB. Dynamic analysis of cytosolic glucose and ATP levels in yeast using optical sensors. The Biochemical journal 432, 399–406 (2010).

37. Cabantous S, Waldo GS. In vivo and in vitro protein solubility assays using split GFP. Nature methods 3, 845–854 (2006).

38. Bamber L, Harding M, Monné M, Slotboom DJ, Kunji ER. The yeast mitochondrial ADP/ATP carrier functions as a monomer in mitochondrial membranes. Proceedings of the National Academy of Sciences of the United States of America 104, 10830–10834 (2007).

39. Duncan AL, Ruprecht JJ, Kunji ERS, Robinson AJ. Cardiolipin dynamics and binding to conserved residues in the mitochondrial ADP/ATP carrier. Biochimica et biophysica acta Biomembranes 1860, 1035–1045 (2018).

40. Klingenberg M. The ADP and ATP transport in mitochondria and its carrier. Biochimica et biophysica acta 1778, 1978–2021 (2008).

41. van Roermund CW, Schroers MG, Wiese J, Facchinelli F, Kurz S. The Peroxisomal NAD Carrier from Arabidopsis Imports NAD in Exchange with AMP. 171, 2127–2139 (2016).

42. van Roermund CW, Ijlst L, Wagemans T, Wanders RJ, Waterham HR. A role for the human peroxisomal half-transporter ABCD3 in the oxidation of dicarboxylic acids. Biochimica et biophysica acta 1841, 563–568 (2014).

43. Abraham EH, et al. The multidrug resistance (mdr1) gene product functions as an ATP channel. Proceedings of the National Academy of Sciences of the United States of America 90, 312–316 (1993).

44. Reisin IL, et al. The cystic fibrosis transmembrane conductance regulator is a dual ATP and chloride channel. The Journal of biological chemistry 269, 20584–20591 (1994).

45. Roman RM, et al. Hepatocellular ATP-binding cassette protein expression enhances ATP release and autocrine regulation of cell volume. The Journal of biological chemistry 272, 21970–21976 (1997).

46. Jansen RS, et al. ABCC6-mediated ATP secretion by the liver is the main source of the mineralization inhibitor inorganic pyrophosphate in the systemic circulation-brief report. Arterioscler Thromb Vasc Biol 34, 1985–1989 (2014).

47. Costello JL, Passmore JB, Islinger M, Schrader M. Multi-localized Proteins: The Peroxisome-Mitochondria Connection. Sub-cellular biochemistry 89, 383–415 (2018).

48. Cartwright JL, Gasmi L, Spiller DG, McLennan AG. The Saccharomyces cerevisiae PCD1 gene encodes a peroxisomal nudix hydrolase active toward coenzyme A and its derivatives. The Journal of biological chemistry 275, 32925–32930 (2000).

49. Xu W, Dunn CA, Bessman MJ. Cloning and characterization of the NADH pyrophosphatases from Caenorhabditis elegans and Saccharomyces cerevisiae, members of a Nudix hydrolase subfamily. Biochemical and biophysical research communications 273, 753–758 (2000).

50. Stjepanovic G, Baskaran S, Lin MG, Hurley JH. Vps34 Kinase Domain Dynamics Regulate the Autophagic PI 3-Kinase Complex. Molecular cell 67, 528–534.e523 (2017).

51. Bartoszewska M, et al. Peroxisomal proteostasis involves a Lon family protein that functions as protease and chaperone. The Journal of biological chemistry 287, 27380–27395 (2012).

52. Pomatto LC, Raynes R, Davies KJ. The peroxisomal Lon protease LonP2 in aging and disease: functions and comparisons with mitochondrial Lon protease LonP1. Biological reviews of the Cambridge Philosophical Society 92, 739–753 (2017).

